# The anti-inflammatory peptide Catestatin blocks chemotaxis

**DOI:** 10.1101/2020.11.23.393934

**Authors:** Elke M. Muntjewerff, Kristel Parv, Sushil K. Mahata, Mia Phillipson, Gustaf Christoffersson, Geert van den Bogaart

## Abstract

Increased levels of the anti-inflammatory peptide catestatin (CST), a cleavage product of the pro-hormone chromogranin A, correlates with less severe outcomes in hypertension, colitis and diabetes. However, it is unknown how CST reduces the infiltration of monocytes and macrophages in inflamed tissues. Here, we report that CST blocks leukocyte migration towards inflammatory chemokines. By *in vitro* and *in vivo* migration assays, we show that although CST itself is weakly chemotactic, it blocks migration of monocytes and granulocytes to inflammatory attracting factor CC-chemokine ligand 2 (CCL2) and macrophage inflammatory protein 2 (MIP-2). Moreover, it directs CX_3_CR1^+^ macrophages away from pancreatic islets. These findings support the emerging notion that CST is a key anti-inflammatory modulator.

## 1. Introduction

As an immunological response to inflammation, monocytes, granulocytes and leukocytes are attracted to inflamed tissues by chemokines such as CC-chemokine ligand 2 (CCL2, a.k.a. MCP-1) and macrophage inflammatory protein 2 (MIP-2, a.k.a. CXCL2) (1). However, to avoid an excessive response, leukocyte infiltration should be halted for resolution of inflammation, but the mechanisms that govern this are unknown (2). Here, we addressed the potential chemotactic effect of chromogranin A (CgA)-derived peptide Catestatin (CST: hCg_A352-372_) (3). While CST circulates at low nM range, the local concentrations were detected in the μM range in mouse tissues (3-6).

Being an anti-inflammatory peptide, CST reduces inflammation in cardiac and chronic inflammatory diseases (3,7-9). Despite the chemotactic effects of CST (7,10,11), administration of exogenous CST reduces monocyte and macrophage infiltration in the liver, heart and gut in mouse models of type II diabetes, hypertension, atherosclerosis and colitis (4,7,8,12,13). In a colitis model, CST also reduced granulocyte infiltration in the colon (8). In line with this, the adrenal gland, heart, and gut of CST knockout mice display increased macrophage infiltration (4,7,12). In this study, we show that while CST itself is weakly chemotactic, it blocks the extravasation and migration of phagocytes both *in vitro* and *in vivo*. Thus, the anti-inflammatory effects of CST are partly the result of redirecting monocytes and granulocytes away from the inflammation sites.

## 2. Methods; experimental procedures

### 2.1 Animals and human bloods samples

Male and female C57BL/6J (Taconic, Denmark) and *Cx_3_cr1^GFP^* (14) mice weighing 20-26 g were used. All animal experiments were approved by the Regional Animal Ethics committee in Uppsala, Sweden. The research with human blood samples at the Department of Tumor Immunology complies with all institutional and national ethics regulations and has been approved by the ethics committee of Sanquin blood bank. All blood donors were informed of the research and have granted their consent.

### 2.2 Gradientech assay

A CellDirector 2D device (Gradientech) was coated with bovine serum overnight. Human peripheral blood monocytes were isolated from buffy coats of healthy donors as described (15), followed by human microbead CD14^+^ isolation of monocytes according to manufactures’ instructions (130-050-201, Milteny Biotec). Monocytes were activated with LPS for 1 h, washed with PBS, and seeded in the device in 200 μl RPMI-1640 medium. After one hour at 37°C, the two supplied syringes with 1 ml of RPMI-1640 medium, with one containing 5 μM CST were attached to the CellDirector and a flow rate of 5 μl/min was applied. Monocyte movement was visualized with an Axiovert 200 M microscope with a 5x objective (Zeiss, Jena, Germany). Movies were recorded at 2 frames/min for 3 hours. Cell movement was analysed using the Tracking Tool PRO software (Gradientech). 0.5 nM CCL2 (300-04, PeproTech) was used as a positive control.

### 2.3 Cremaster muscle imaging

Monocyte and granulocyte (Ly6G-mAb) migration was imaged in the cremaster muscle of mice superfused with pre-warmed (37°C) bicarbonate-buffered saline solution (pH 7.4) (16) containing CST (5 μM) and/or MIP-2/CXCL2 (0.5 nM) (250-15, PeproTech) was used as a positive control. A bright-field intravital microscope (Leica DM5000B) with a 25×/0.6W (Leica) objective and connected to an Orca R2 camera (Hamamatsu; Volocity acquisition software) was used to record movies of five minutes at 0, 30, 60, 90 min after cytokine addition. Venules with diameter range of 20-30 μm were imaged. Movies were analysed using ImageJ and corrected using the Hyperstackreg ImageJ macro. For rolling flux, all cells rolling in the vessel were counted. For rolling speed, velocity over a 100 μm section of the vessel was analysed. In the same 100 μm section, cells were considered adherent if they remained stationary for at least 3 min.

### 2.4 Aortic ring assay with pancreatic islet culture

Aortic ring isolation was carried out as previously described (17). Briefly, 13-16-week-old *Cx_3_cr1^GFP^mice* were euthanized, followed by dissection of the thoracic aorta. Under a stereo-microscope, extraneous fat, tissue, and branching vessels were carefully removed, and perfused with serum-free OptiMEM medium (Thermo Fisher) with penicillin-streptomycin solution. The aorta was sectioned into 1 mm thick rings. After overnight starvation in serum-free Opti-MEM medium, rings were embedded in 1 mg/ml rat tail collagen I (#ALX-522-435-0100, Enzo Life sciences) adjacent to pancreatic islets (2-5 islets per ring), which were isolated from C57BL/6 mice as described before (18), in 8 well Nunc Lab-Tek II microscope chambers (Thermo Fisher). After 1 h, embedded rings were cultured with 300 μl of OptiMEM with 2.5% FBS, 11.1 mM glucose, penicillin-streptomycin, M-CSF (40 ng/ml) to stimulate CX_3_CR1^GFP+^ macrophage survival and 5 μM CST for six days. On day six, rings were imaged using a Zeiss LSM700 (Carl Zeiss) confocal microscope. The numbers of CX_3_CR1^GFP+^ cells were quantified using the image analysis software Imaris (Bitplane). The location of the CX_3_^GFP+^ cells was determined using the Surface Center of Mass Position to Spots object plugin after manually defining the aorta. For analyzing angiogenesis, staining with anti-CD31 antibody conjugated to Alexa Fluor 647 (#102515, Biolegend) was carried out prior to imaging. Aortic rings that did not show any sprouting were excluded from further analysis. Vessels were analyzed using Fiji image analysis software (19). Sprouts that originated directly from the ring endothelium were considered main sprouts, and branches as divarications from main sprouts.

### 2.5 Statistical data analysis

Data are expressed as mean ± SEM. One-way ANOVA with Bonferroni post-hoc tests or non-parametric Mann-Whitney test were applied for multiple comparisons. Outliers were identified using ROUT test (Q=1%). A value of p < 0.05 was considered statistically significant.

## 3. Results & Discussion

Although human blood monocytes migrated towards a high (but physiological) concentration of CST (5 μM), this was less efficient compared to the canonical inflammatory chemokine CCL2 (0.5 nM) (Fig. 1A-C), reinforcing a weak chemoattractive effect of CST (7,10,11). To confirm this *in vivo*, we performed imaging of the cremaster muscle (Fig. 1D) (16). Upon perfusion of the muscle with CST (5 μM), phagocytes (monocytes and granulocytes) decreased their speed and attached to the vessel wall with similar efficiency as of the inflammatory chemotactic agent MIP-2 (0.5 nM) (Fig. 1D-F, Fig. S1). Thus, both our *in vivo* and *in vitro* migration assays show that CST is weakly chemotactic, raising the question how CST can reduce monocyte and granulocyte infiltration in inflamed tissues such as the liver (diet induced obese mice), intestine (colitis model), heart (hypertension model) and atheromatous plaques (atherosclerosis model) (4,7,8,12,13).

**Fig. 1:**
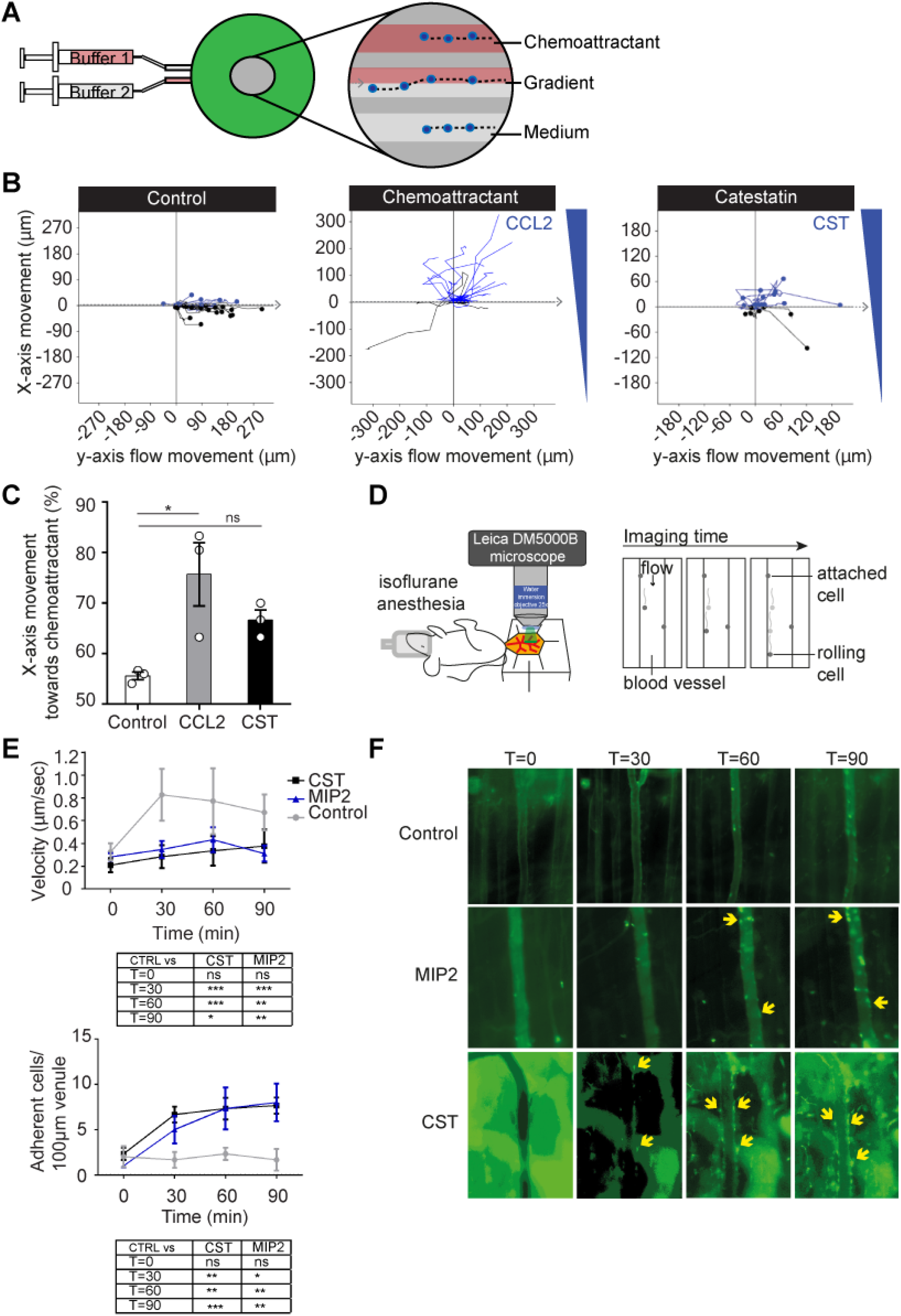
CST is weakly chemotactic. **(A)** Scheme showing set-up of Gradientech migration assay. Two syringes filled with buffer +/- chemoattractant were connected to the device (green) to create a flow (*x*-direction) and perpendicular (*y*) cytokine gradient. The inset shows migration of monocytes along the flow and towards the chemoattractant. **(B)** Representative tracks of human monocytes showing the *x*- and *y*-movement of individual cells upon exposure to the indicated buffer, 5 μM CST or 0.5 nM CCL2. **(C)** Quantification of panel B (N=3). **(D)** Scheme showing set-up of cremaster muscle imaging in mice to visualize phagocyte (monocytes and granulocytes) extravasation *in vivo*. **(E)** Phagocyte rolling velocity (top) and attachment (bottom) upon overflowing the muscle with buffer (control, gray), 0.5 nM MIP-2 (blue) or 5 μM CST (black) (N=3, two-way ANOVA). **(F)** Representative images of granulocyte attachment as visualized by Ly6G-mAb (green) to the vessel wall upon only buffer, MIP-2 or CST stimulation. *: P<0.05; **: P<0.01; ***P<0.001; ns: not significant.

To address how CST affects macrophage chemotaxis to inflamed tissues, we used the aortic ring vessel model (17) (Fig. 2A), which is based on the co-embedding of part of the aorta of *Cx_3_cr1^+/gfp^* transgenic mice adjacent to isolated pancreatic islets (20). These islets secrete chemokines, such as vascular endothelial growth factor (VEGF)-A, resulting in the directional macrophage migration from the aortic ring as well as vessel growth towards the pancreatic islets. Migration of CX_3_CR1^+^macrophages from the aortic ring was visualized by fluorescence microscopy (19) (Fig. 2B, S2). As expected, the CX_3_CR1-macrophages moved towards the pancreatic islets in absence of CST (Fig 2B). However, perfusing the aortic ring with CST (5 μM) resulted in a lower number of CX_3_CR1^+GFP^ macrophages migrating towards the pancreatic islets (Fig 2B), indicating that CST blocked directional migration. Interestingly, we also observed that CST is pro-angiogenic, as it increased both the amount and length of the sprouts and branches emanating from the aortic rings (Fig. 2C-D, S3).

**Fig. 2:**
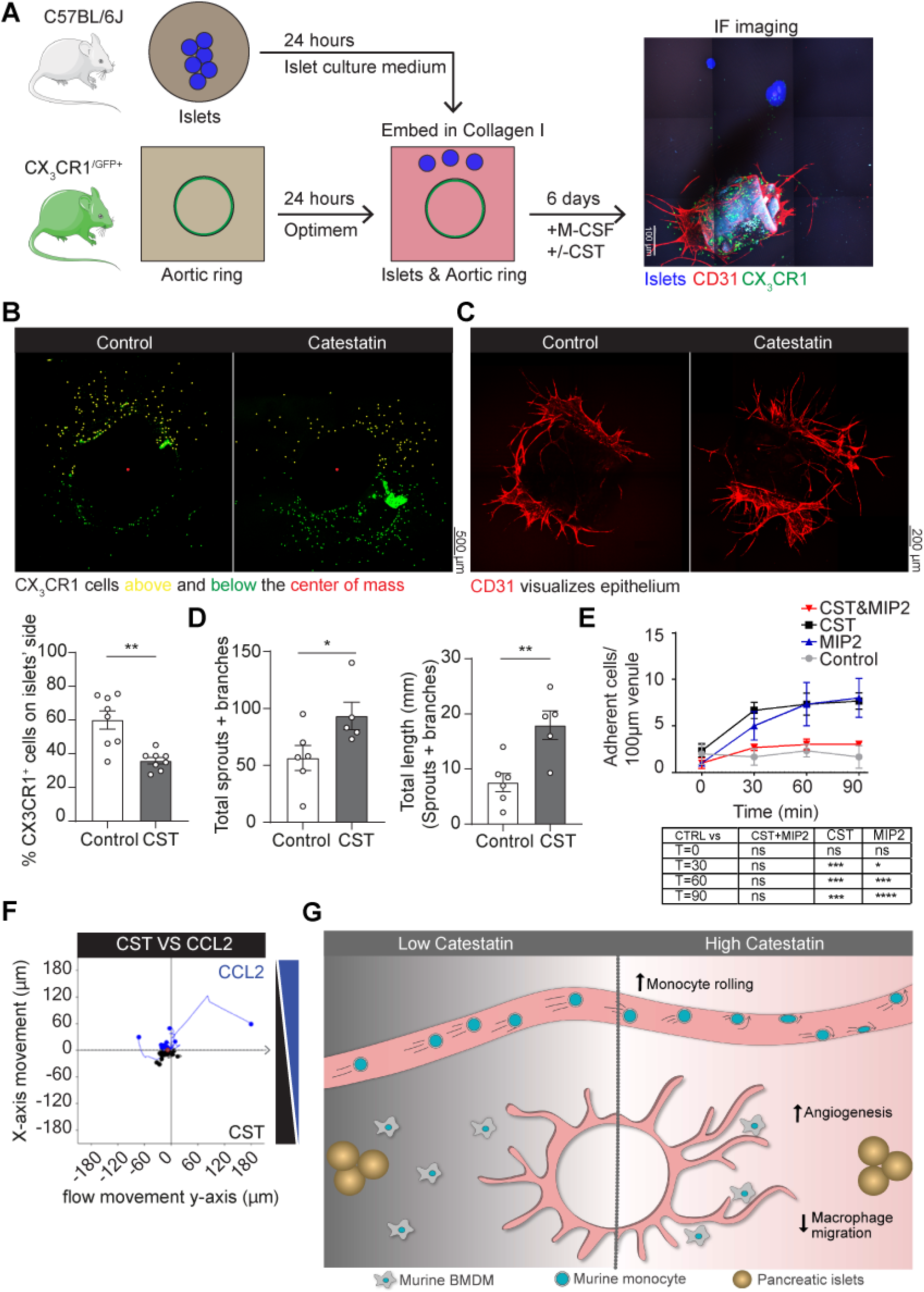
CST blocks migration induced by inflammatory chemokines and promotes angiogenesis. **(A)** Scheme showing set-up of aortic ring assay. Aortic ring was isolated from CX_3_CR1-GFP mice and embedded adjacent to pancreatic islets in collagen I. Image shows islets (blue), CD31 (red) and CX_3_CR1 (green). **(B)** Representative images of CX3CR1-macrophage migration upon control or CST stimulation of the aortic ring. The graph shows the percentage of cells above (yellow) the center of mass (N=8). **(C)** Representative images of vessels by CD31 (red) upon control or CST stimulation of the aortic ring. **(D)** Quantification of angiogenesis. Total number of sprouts and branches (left) and their length (right) (N=5-6). **(E)** Cremaster muscle imaging. Phagocyte attachment to vessel wall upon overflowing the muscle with buffer (control, gray) and buffer with the chemoattractant MIP-2 (blue), CST (black) or both (red) (N=3, two-way ANOVA). **(F)** Gradientech migration assay. Representative *x*- and *y*-movement of human monocytes exposed to opposite gradients of CST and CCL2 (N=3). **(G)** Model showing leukocyte extravasation in presence of low and high concentrations of CST. Mann-Whitney test *: P<0.05; **: P<0.01; ***P<0.001; ****P<0.0001; ns: not significant.

The loss of directional cell migration to the pancreatic islets might be caused by blockage of chemokine-induced cell migration by CST. To investigate this possibility, we performed intravital imaging of the cremaster muscle, but this time for CST in combination with MIP-2. This resulted in the inverse effect compared to CST or MIP-2 alone: release of attached cells from the vessel wall and reduced migration of cells into the tissue (Fig. 2E, S4), indicating that despite being weakly chemotactic, CST blocks MIP-2 elicited phagocyte recruitment. To further confirm this, we performed an *in vitro* migration assay, where human monocytes were stimulated with a gradient of CCL2 in presence of CST (Fig. 2F). Similar to our findings with the intravital imaging, CST blocked monocyte migration towards the CCL2.

Although CST counteracts the chemoattraction by inflammatory cytokines (Fig. 2G), the question remains open which receptor(s) CST utilize to exert these effects on cell migration. We speculate that this might be a G-protein coupled receptor (GPCR), since GPCRs are actively involved in leukocyte migration (21) coupled with expression of GPCRs in all cell types responsive to catestatin (e.g. monocytes (10), neutrophils (22,23), macrophages (4,7,8,12,13), endothelial (13,24) and mast cells (11)), we speculate that CST might act through this receptor type. We have not only shown how CST reduces the infiltration of monocytes and macrophages in inflamed tissues (4,7,8,12,13), but offer a possible mechanistic explanation for the correlation of CST levels with improved disease outcome in patients suffering from chronic diseases (4-6), reinforcing CST as a therapeutic target for treatment of diseases associated with chronic inflammation.

## 4. Author contributions

E.M.M., G.C., S.K.M. and G.v.d.B. designed the study. E.M.M, K.P., G.C. designed and performed the experiments. E.M.M. and G.v.d.B wrote the manuscript and all authors participated in discussing and editing of the manuscript.

## 5. Disclosure of conflict of interest

The authors declare that the research was conducted in the absence of any commercial or financial relationships that could be construed as a potential conflict of interest.

## 6. Funding

G.v.d.B. is funded by a Young Investigator Grant from the Human Frontier Science Program (HFSP; RGY0080/2018), and a Vidi grant from the Netherlands Organization for Scientific Research (NWO-ALW VIDI 864.14.001). G.v.d.B has received funding from the European Research Council (ERC) under the European Union’s Horizon 2020 research and innovation programme (grant agreement No. 862137. S.K.M. is supported by a grant from the US Department of Veterans Affairs (I01BX000323). G.C. is supported by grants from the Swedish Research Council and the Swedish Society for Medical Research. E.M.M is supported by a short-term EMBO fellowship (EMBO7887).

**Sup. 1: Attachment of granulocytes and monocytes to vessel wall. (A)** Venules of the cremaster muscle were overflown with bicarbonate-buffered saline buffer (buffer only control), the chemoattractant MIP-2 or CST as shown in main Fig. 1D-F. Graph shows quantification of rolling cells (cells/min). **(B)** Quantification of cell in tissue. **(C)** Representative brightfield snapshots of *in vivo* cremaster muscle imaging as in main figure 1D-F. **(C)** Quantification of adherent granulocytes (visualized by Ly6G-mAb, main Fig. 1F) and monocytes (brightfield, panel C) after 0, 30, 60 and 90 minutes (N=1-2).

**Sup. 2: Quantification of CX3XR1+ cell movement in the aortic ring model. (A)** Brightfield image of the aortic ring with islets. **(B)** Description of CX3CR1-cell movement quantification by determination of total amount outside the aortic ring (endothelium), center of mass (red spot) and the islet side (black arrow).

**Sup. 3: Branches and sprouts in the aortic ring assay. (A)** Representative images of angiogenesis quantification of main figure 2C-D. The images show the ring endothelium, main sprout (red), branch (gray) **(B)** Quantification of total number of sprouts and branches separately and their length (N=5-6). Mann-Whitney test *: P<0.05; ***P<0.001; ns: not significant.

**Sup. 4: The combination of CST and MIP-2 reduced chemotaxis.** Venules of the cremaster muscle were overflown with bicarbonate-buffered saline buffer (buffer only control), the chemoattractant MIP-2 or CST, as shown in main Fig. 1D. Graph shows quantification of tissue migration **(A)**, rolling cells (cells/min) **(B)** and velocity **(C)** upon CST, MIP-2 or stimulation with both (N=3, two-way ANOVA) *: P<0.05; **: P<0.01; ***P<0.001; ****P<0.0001; ns: not significant.

## Notes

### Competing Interest Statement

The authors have declared no competing interest.

